# REGALS: a general method to deconvolve X-ray scattering data from evolving mixtures

**DOI:** 10.1101/2020.12.06.413997

**Authors:** Steve P. Meisburger, Da Xu, Nozomi Ando

**Author notes:** These authors contributed equally (Steve P. Meisburger, Da Xu).

## Abstract

Mixtures of biological macromolecules are inherently difficult to study using structural methods, as increasing complexity presents new challenges for data analysis. Recently, there has been growing interest in studying evolving mixtures using small-angle X-ray scattering (SAXS) in conjunction with time-resolved, high-throughput, or chromatography-coupled setups. Deconvolution and interpretation of the resulting datasets, however, are nontrivial when neither the scattering components nor the way in which they evolve are known a priori. To address this issue, we introduce the REGALS method (REGularized Alternating Least Squares), which incorporates simple expectations about the data as prior knowledge and utilizes parameterization and regularization to provide robust deconvolution solutions. The restraints used by REGALS are general properties such as smoothness of profiles and maximum dimensions of species, which makes it well-suited for exploring datasets with unknown species. Here we apply REGALS to analyze experimental data from four types of SAXS experiment: anion-exchange (AEX) coupled SAXS, ligand titration, time-resolved mixing, and time-resolved temperature jump. Based on its performance with these challenging datasets, we anticipate that REGALS will be a valuable addition to the SAXS analysis toolkit and enable new experiments. The software is implemented in both MATLAB and python and is available freely as an open-source software package.

## 1 Introduction

Small angle X-ray scattering (SAXS) is a widely used technique for obtaining structural information from macromolecules in solution [1]. Increasingly, SAXS is applied to evolving mixtures of different molecules or conformational states [2, 3] during titrations [4], chromatographic separation [5], or time-resolved experiments [6–8]. However, because of the fundamental limitations in the information content of the SAXS signal [9], multiple structures in a mixture cannot be resolved from each profile in an unambiguous manner. This inherent ambiguity can be mitigated by combining multiple measurements and carefully incorporating prior knowledge. The individual components can then be separated mathematically by analyzing the dataset as a whole using a physicochemical model for how the mixture evolves [10–12] or known scattering curves of each component [13]. Often, however, both the scattering curves and physicochemical model are unknown before the experiment is performed and must be inferred from the data itself. In such cases, the challenge is to identify appropriate mathematical tools to incorporate more general, physically motivated restraints that lead to a reliable and accurate model-free separation.

In dilute solution, SAXS intensities from non-interacting components combine linearly in proportion to their relative concentrations. A SAXS dataset from a mixture can therefore be described as the convolution of the concentration and SAXS profiles, and deconvolution can be performed using matrix factorization techniques such as singular value decomposition (SVD) [14, 15]. However, to recover the scattering from each component, the basis vectors from SVD must be recombined using prior knowledge about what constitutes a physically valid solution. The field of chemometrics has developed a number of algorithms for solving this problem, known as multivariate curve resolution or MCR [16, 17]. When a physicochemical model is available, the alternating least squares (MCR-ALS) algorithm can perform deconvolution using the model as a hard restraint [17]. In the context of SAXS, deconvolution with hard restraints has been applied to time-resolved experiments [11, 18–20], equilibrium titrations [10, 12, 21, 22], unfolding experiments [23, 24], protein-micelle interactions [25], and fibril formation [26]. Interestingly, MCR can be performed without assuming a hard model by imposing soft restraints such as positivity, unimodality, and local rank [17]. Such model-free deconvolution is seldom applied to SAXS data because soft restraints are rarely sufficient to provide a robust and unique solution on their own [16]. One exception is SAXS data collected with in-line size-exclusion chromatography (SEC-SAXS), where MCR-ALS has been combined with evolving factor analysis (EFA) [27] to separate overlapping elution peaks [28, 29].

Although SVD and MCR algorithms are well suited to certain SAXS experiments, they are a poor fit for other more challenging datasets. A notable example is SAXS data collected with in-line anion exchange chromatography (AEX) [30]. AEX separates according to charge by applying the sample to cationic media and eluting with a salt gradient. In SAXS, the salt gradient produces a changing background scattering that must be accounted for. Because this changing background violates certain assumptions of the EFA method, model-free deconvolution of AEX-SAXS data is not possible with EFA. We previously encountered this issue when analyzing AEX-SAXS data from the large subunit of B. subtilis ribonucleotide reductase (BsRNR) [31]. To overcome this challenge, we incorporated a simple assumption as additional prior information: namely, that the background scattering must change gradually over time. Using the ALS algorithm with smoothness regularization applied to the concentration of background scattering components, we achieved a clean separation of multiple protein and buffer components [31].

Here, we examine the generality of this strategy for the model-free deconvolution of other complex types of SAXS data where traditional “soft” restraints are insufficient. We describe the REGALS (REGularized ALS) toolset and demonstrate its application to a wide variety of SAXS experiments from evolving mixtures. Unlike most deconvolution methods that impose a physicochemical model, REGALS relies on very general parametric models for the SAXS profiles and concentration curves. The models include two types of restraint: smoothness and compact support. In AEX-SAXS, for example, each elution peak is assumed to be non-zero over a particular range (compact support), and the background components are assumed to be smooth. For the BsRNR dataset, we find this is sufficient to deconvolve the protein scattering peaks. In other cases, such as equilibrium titration and time-resolved SAXS, where concentrations are typically non-zero in all (or nearly all) data frames, the assumption that concentrations have compact support is insufficient. However, compact support can be applied to the SAXS profiles in real space by imposing a maximum particle dimension. We show that compact support in real space, as well as boundary conditions applied to the concentration basis functions, provide sufficient information for successful deconvolution of such data. Finally, we introduce the REGALS software package, which is adaptable by design, freely available, and open source.

## 2 Theory

### Background

A dilute, evolving mixture of *K* components scatters X-rays according to the following linear model:

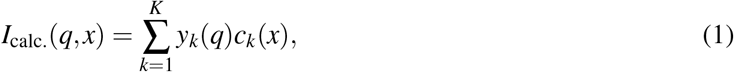

where *y_k_*(*q*) are the individual SAXS profiles and *c_k_*(*x*) are the relative concentrations. The SAXS profiles depend on the scattering vector magnitude *q* = (4*π/λ*) sin *θ*, where *λ* is the X-ray wavelength and 2*θ* is the scattering angle. The concentration profiles depend on an independent variable *x* (representing time, ligand concentration, etc). Since intensities are measured at discrete values of *q* and *x*, Equation 1 can be written in matrix form as follows:

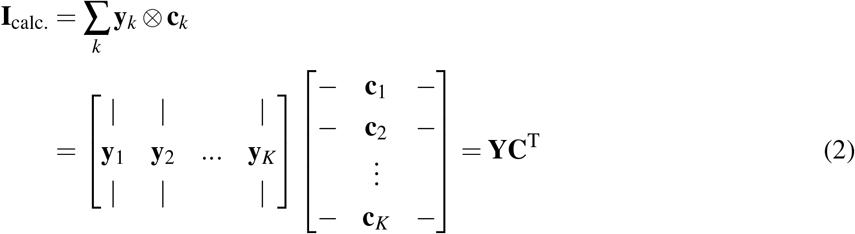

where **I**_calc_. contains scattering profiles arranged side-by-side as column vectors. Here and throughout this section, the intensity matrix has dimensions of *M × N* (*N* scattering profiles with *M* discrete values of *q*). Hence, **Y** is *M* × *K* and **C** is *N* × *K*.

Our aim is to determine **Y** and **C** given the measured intensity **I**_meas_., which contains noise. This is accomplished by minimizing the least-squares error between data and model:

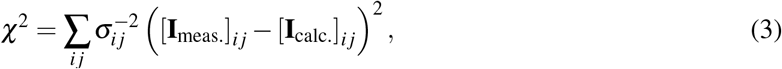

where *σ_ij_* are the standard errors of the measured intensity. In the following, we assume that the experimental errors depend only on *q*, so that Equation 3 can be written as a Frobenius norm of the error-weighted residual:

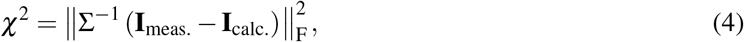

where **Σ** is a diagonal matrix with 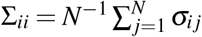. This simplifying assumption is approximately correct for the datasets considered here.

In general, minimizing *χ*^2^ is not sufficient to determine **Y** and **C** uniquely. The main issue is that basis vectors can be mixed (or “rotated”) without changing *χ*^2^: for any non-singular *K × K* matrix **Ω**, replacing **Y** → **YΩ** and **C** → **CΩ**^−T^ leaves the product **YC**^T^ unchanged. Thus, the primary challenge of deconvolution is to impose appropriate restraints that provide a unique and physically meaningful solution.

Deconvolution problems resembling Equation 2 arise in many experimental contexts. A common approach is to apply SVD [14, 15], by which an error-weighted data matrix is decomposed as follows:

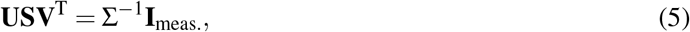

where **U** has the left singular vectors as columns, **V** contains the right singular vectors as columns, and **S** contains the singular values along the diagonal in decreasing order. The uniqueness of the decomposition results from the fact that the singular vectors are an orthonormal basis.

The singular values *s_j_* = *S_jj_* are positive and indicate the importance, or weight, for each pair of left and right singular vectors. When the number of observations (*N*) is much larger than the number of independent components in the signal (which is generally the case for examples studied here), most of the singular values will be small and represent the noise in the data, while a few large singular values correspond to the signal of interest. To detect significant singular values, it is useful to calculate a normalized singular value, as follows:

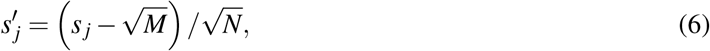

where *M* and *N* are the number of rows and columns of the data matrix. If no signal is present, random matrix theory shows that 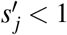 in the limit where the data matrix is large (see [32] and references therein). Thus, components corresponding to signal above the noise are expected to have 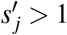.

By retaining only the *K* most important singular vectors (**U** → **U**_*K*_, etc.), one obtains an approximate (reduced rank) representation of the data. Thus, a solution for **Y** and **C** can be constructed from SVD as follows:

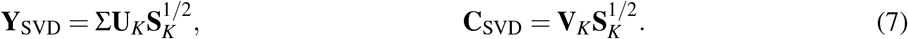

Here, the singular value weights have been distributed evenly between the SAXS and concentration basis vectors, but other choices could be made depending on the normalization conditions.

Although SVD provides a unique low-rank decomposition of the data, the orthonormality of the singular vectors often produces non-physical results. For instance, the component SAXS profiles or concentrations might have negative values. It is therefore often necessary to further unmix (or “rotate”) the SVD basis vectors by applying physical restraints [10, 19, 23, 25]. In traditional MCR techniques, physical restraints are imposed using “hard” or “soft” models, whose applicability depends on the type of experiment performed and prior knowledge. Alternatively, prior information can be imposed through Tikhonov-Miller regularization, where additional functions are minimized at the same time as *χ*^2^ [33, 34]. As described above, in an AEX-SAXS experiment, the expectation that background scattering varies gradually over time can be enforced using a regularization function that penalizes large oscillations [31].

Regularization is also used in conventional SAXS data analysis to infer the pair distance distribution function, or *P*(*r*), from the measured intensity [35]. Essentially, *P*(*r*) represents the probability of two electrons having a distance r in the sample, and it is related to the scattering intensity by a Fourier transform:

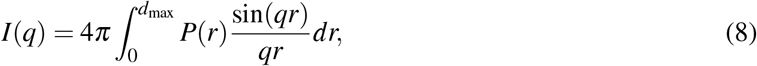

where the integral terminates at the maximum particle dimension, *d*_max_(since *P*(*r*) = 0 for *r* > *d*_max_). Although Equation 8 can be inverted analytically, in practice the intensity is measured over a finite q-range, and thus, inversion is an ill-posed problem. Since the Fourier transform is a linear operator, Tikhonov-Miller regularization can be applied. *P*(*r*) is discretized as a vector **u** of length *R*, which samples values of *P*(*r*) on a uniform grid with spacing Δ*r*. Equation 8 can then be written as

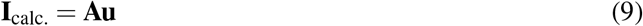

where **I**_calc_ is a vector of length *M*, and **A** is an *M × R* matrix with elements

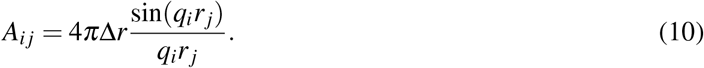

The standard indirect Fourier transform (IFT) method for SAXS data minimizes the *χ*^2^ between I_calc_. and I_meas_. plus a regularization term:

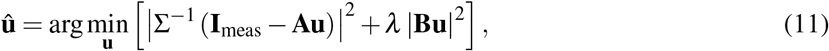

Typically, the matrix **B** performs a discrete approximation of the second derivative [36, 37], which enforces smoothness by penalizing wildly oscillating solutions. The regularization parameter (or Lagrange multiplier) λ controls the tradeoff between minimizing *χ*^2^ and minimizing the regularization function. The optimization problem is solved by the method of normal equations, with the (formal) result:

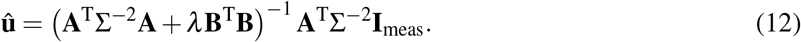

In this study, we describe a general method for deconvolving SAXS data from mixtures that applies regularization to both the concentration and SAXS profile basis vectors. We first formulate the deconvolution problem (Equation 2) using a parametric representation of the basis vectors, similar to the IFT example above. This parametric form allows the SAXS profiles to be represented in the real-space (P(r)) basis if desired. Then, we describe the REGALS algorithm for minimizing the sum of *χ*^2^ (Equation 3) and regularization terms.

### 2.2 Deconvolution by regularized least squares

In order to deconvolve SAXS data from evolving mixtures, we introduce a method to impose mathematical constraints that embody prior information (or general expectations) about a SAXS experiment. The first way that constraints are imposed is through a parameterization of the basis vectors:

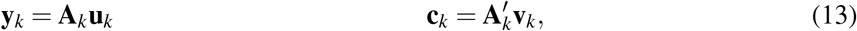

where **u**_*k*_ and **v**_*k*_ are the parameter vectors for the SAXS profile and concentration bases respectively. Here and in the following equations, primed functions or matrices refer to the concentration basis, in order to distinguish them from the SAXS profile basis. We implemented three types of basis vector: simple, smooth, and real-space (Figure 1a). In a simple basis vector, **A**_*k*_ is the identity matrix and the parameter vector encodes the basis vector directly. In a smooth basis vector, **A**_*k*_ performs a linear interpolation from a uniform grid of control points to the experimental grid, which need not be uniform. Finally, in a real-space basis vector (which applies exclusively to SAXS profiles), **u**_*k*_ samples *P*(*r*) on a uniform grid and **A**_*k*_ is given by Equation 10. Crucially, the global model can contain a mixture of different parameterizations. This model was implemented using a flexible object hierarchy (Figure 1b) as described in Methods.

**Figure 1:**
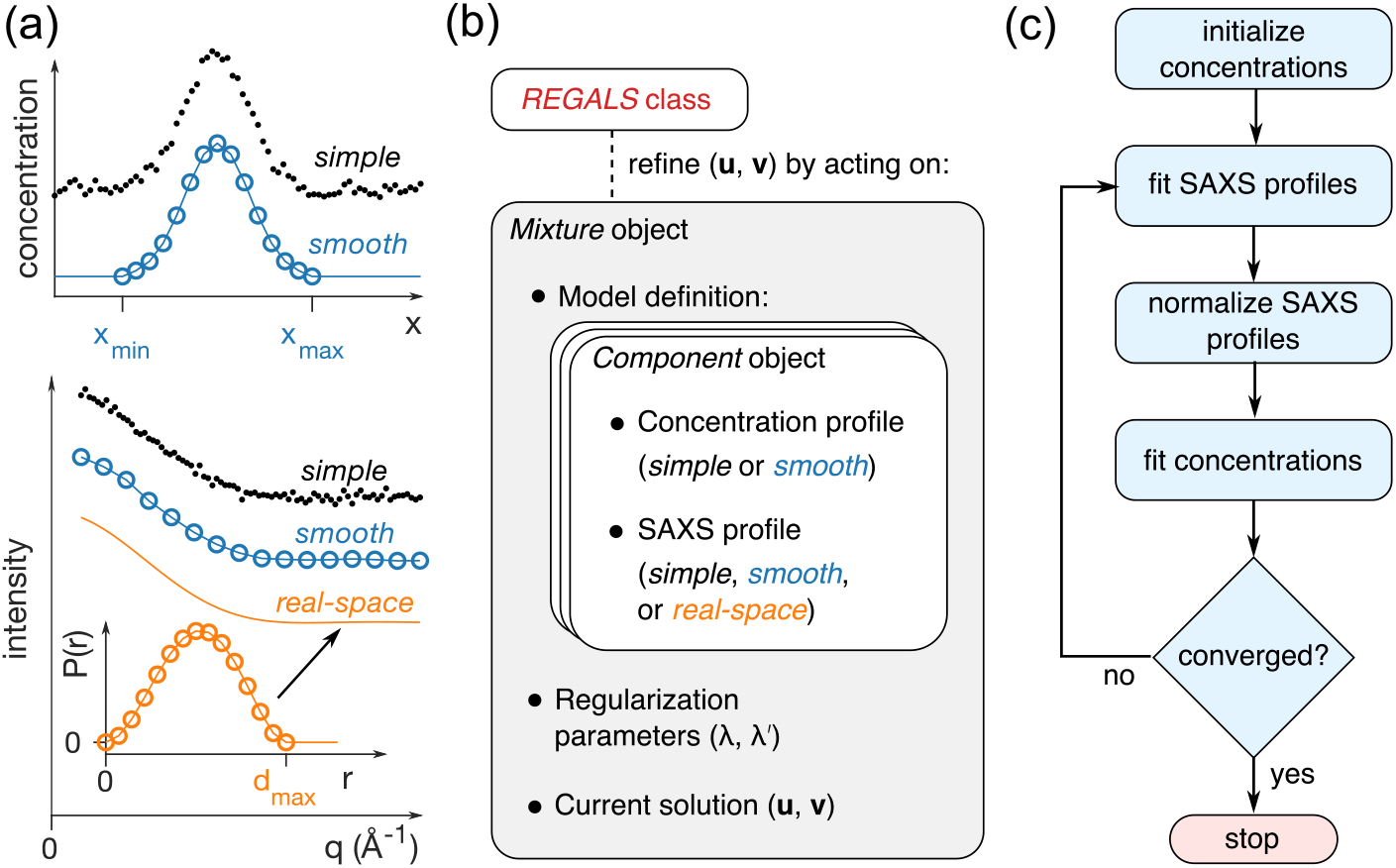
Overview of the REGALS method. (**a**) Parametric basis vectors representing concentrations (top panel) and SAXS profiles (bottom panel). In simple vectors, each sample (*q* or *x*) is given an independent parameter (black dots). A smooth vector represents the data by linear interpolation between control points (blue circles) over the region of support (*x*_min_ and *x*_max_, top panel). A real-space vector samples the *P*(*r*) function (orange circles) up to the maximum particle dimension (*d*_max_), and corresponding SAXS intensities (orange curve) are calculated by Fourier transform (Equation 8). (**b**) Experimental restraints are expressed in software by mixing and matching basis vector types using a flexible object heirarchy. The basis vectors representing SAXS profiles (**u**) and concentrations (**v**) are refined by methods in the high-level REGALS class. (**c**) Refinement algorithm based on regularized ALS. At each iteration, regularized linear least-squares fits are performed on the SAXS profiles (Equation 17) and concentrations (Equation 18) in an alternating fashion until a user-specified convergence test is satisfied.

The second way constraints are imposed is through regularization. The regularization functions 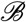 embody prior information (or expectations) about the data, such as smoothness in data or parameter space, and are minimized along with *χ*^2^:

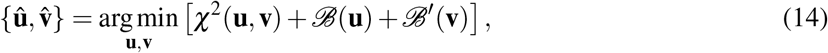

Here, **u** and **v** refer to global parameter vectors that are constructed by concatenating parameter vectors for the individual basis functions (for example, **u** is **u**_1_,…, **u**_*K*_ placed end to end), and *χ*^2^ is calculated from Equations 4 and 13 as follows:

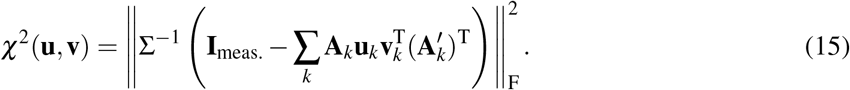

The regularization functions are a sum of quadratic regularizers acting on each component’s parameter vector:

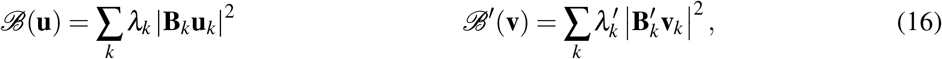

The regularization parameters *λ_k_* and 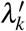 control the tradeoff between minimizing *χ*^2^ and each regularizing function. For smoothness regularization, **B**_*k*_ is a discrete approximation of the second derivative [37]. Zero boundary conditions are optionally imposed by removing the parameters on the boundary and deleting the corresponding rows of **A**_*k*_ and **B**_*k*_ [37].

### 2.3 REGALS algorithm

The optimization problem described in the previous section (Equations 14, 15, and 16) is nonlinear, and therefore does not afford a straightforward solution. We chose to adapt the alternating least squares (ALS) algorithm, which is often used in classic MCR [16, 17, 38]. ALS replaces the single nonlinear optimization problem with two linear problems that are solved in an alternating fashion over many iterations: beginning with an initial guess, one set of basis functions is optimized (e.g. the concentrations) with the other held fixed, and then the other basis functions are optimized. This is repeated until the change in basis vectors from one iteration to the next is smaller than a certain tolerance or the maximum number of iterations has been reached.

The REGALS algorithm solves Equation 14 iteratively using ALS with regularization (Figure 1c). First, an initial guess is made for the concentration basis parameters (**v**). This can be supplied by the user or generated automatically based on the parameterization type and boundary conditions. In the first leastsquares step, the SAXS basis functions are optimized while the concentrations are held fixed:

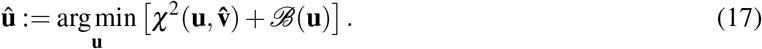

Then, the profiles are normalized according to a their parameterization type; for simple and smooth types, the parameters are divided by the root-mean-squared value, while for the real-space type, parameters are normalized by the scattering intensity at *q* = 0 calculated from the area under the *P*(*r*) curve (see Equation 8). In the second least-squares step, the concentration basis functions are optimized while the SAXS profiles are held fixed:

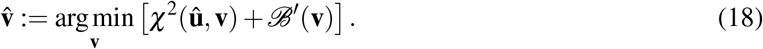

Statistics are calculated at this stage, including the change in the basis vector from the previous iteration (sum of the absolute value of the difference) as well as the *χ*^2^ for the current model (Equation 3). Finally, the cycle is repeated until a reaching convergence according to user-specified termination conditions. Further details about parameter estimation, error analysis, and implementation can be found in Methods.

## 3 Results and Discussion

### 3.1 REGALS deconvolution of AEX-SAXS data

During an AEX separation, sample bound to the column is eluted by flowing buffer with increasing salt concentrations. The main challenge in deconvolving AEX-SAXS data is to account for the changing background scattering from the buffer. We analyzed a dataset previously reported for the large subunit of BsRNR [31], which eluted from the column in two main peaks during a linear gradient of 100 to 400 mM NaCl (Figure 2a). The salt gradient produced a rising background intensity during elution, seen clearly in a plot of the total intensity per frame (Figure 2b, top panel).

**Figure 2:**
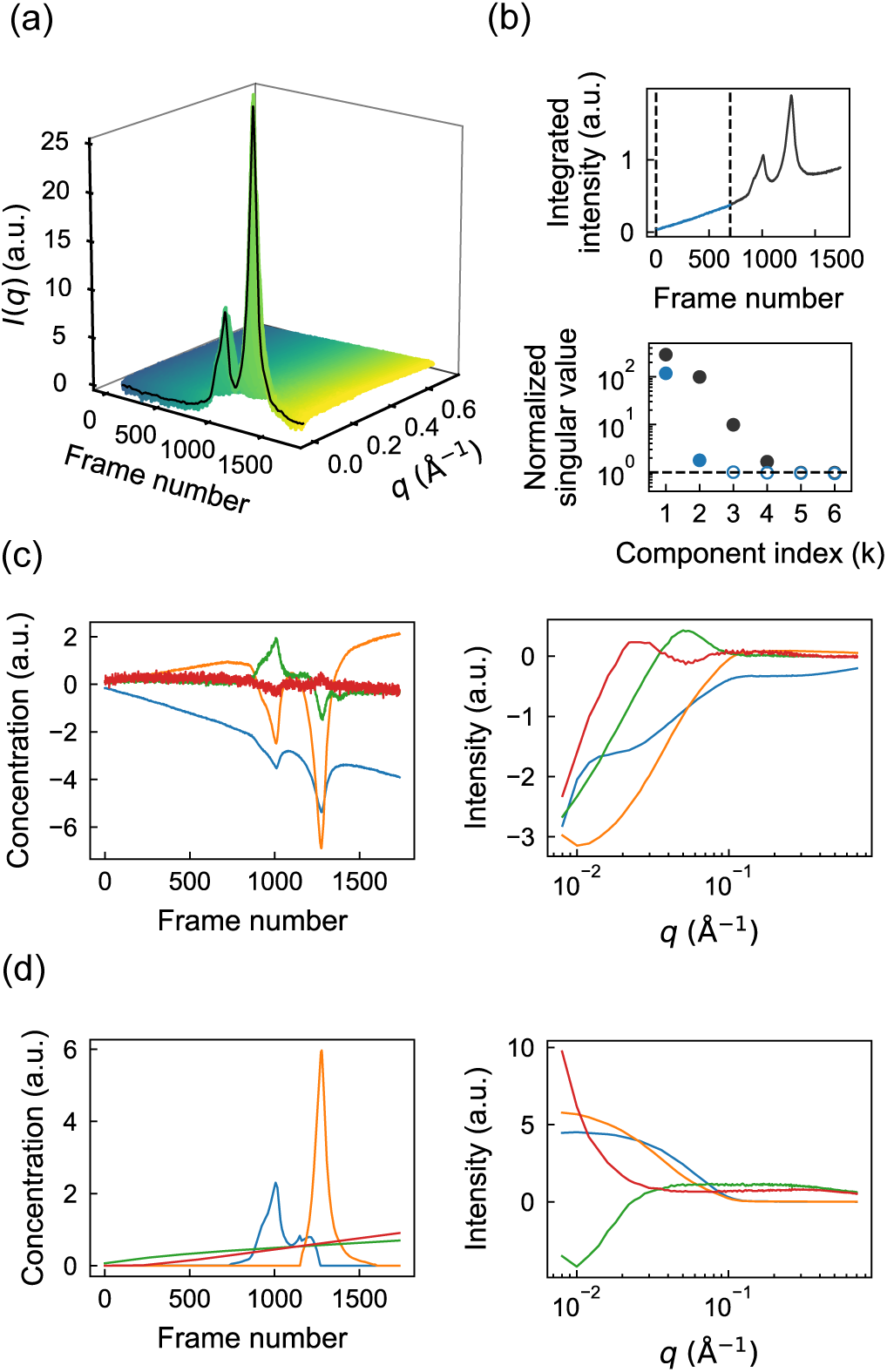
REGALS deconvolution of an AEX-SAXS dataset with a changing background. (**a**) The scattering intensities obtained in a previously reported AEX-SAXS experiment on the large subunit of *Bs*RNR [31] plotted as a function of frame number. (**b**) In the top panel, the integrated intensities across the elution display two prominent peaks over a rising background. SVD of the full dataset shows four significant singular values above the noise level (gray circles above the dashed line in bottom panel, see Equation 6). SVD of only those scattering profiles prior to the protein peaks (blue region between dashed lines in top panel) shows two significant singular values (blue filled circles in bottom panel), indicating the presence of two background components. (**c**) The deconvolution derived from SVD of the full dataset (Equation 7) is non-physical. On the left are the concentration profiles (right singular vectors), and on the right, the corresponding scattering profiles (left singular vectors). (**d**) REGALS gives physical concentration profiles (left panel) and scattering profiles (right panel).

First, we performed SVD to estimate the number of scattering components associated with the protein and background signals. SVD of the entire dataset yields four significant singular values (Figure 2b, bottom panel, black circles). To determine which of these four correspond to buffer vs. protein, we repeated SVD on a truncated dataset consisting of the first 700 frames, collected before the protein elution (Figure 2b, top panel, blue region). Interestingly, this region alone produces two significant singular values (Figure 2b, bottom panel, blue circles), suggesting that two components are needed to describe the background, and that the remaining two correspond to protein. Inspection of the basis vectors obtained from SVD of the full dataset (Equation 7) further confirms this assignment. A rising background signal is present in two of the concentration profiles (Figure 2c, left panel, orange and blue curves). However, it is also evident in the concentration profiles that protein peaks appear in all four components, mixing with the background in two cases, and that the corresponding SAXS profiles (Figure 2c, right panel) are similarly non-physical, containing negative intensities. The fact that many of the concentration and SAXS profiles oscillate around zero is expected given the orthonormality restraint imposed on the SVD basis vectors. Thus, although SVD provides useful information on the number and types of scattering components, different restraints are needed in the deconvolution process to obtain a physically meaningful interpretation.

With initial insight from SVD, we next constructed a Mixture model (Figure 1b) that takes into account basic expectations about the data. The simplest assumption is that each peak in the chromatogram corresponds to a different protein component and that the concentrations of the background components should evolve smoothly over the course of elution. Each protein component (C1 and C2) was thus parameterized using a smooth concentration basis vector with a region of support encompassing each peak. In order to arrive at a unique deconvolution, the two background components must be differentiated in some way within the model. Because SVD revealed that one of the background-containing components is close to zero for the first ~ 200 frames (Figure 2c, left panel, orange curve), we modeled one of the background components (B1) to span the full range of frames, while the other (B2) had a region of support beginning at frame 200 with a zero boundary condition there. We implemented this model again using smooth concentration basis vectors for B1 and B2.

Finally, we refined the model using regularization to enforce smoothness of the background components. The SAXS profiles were not parameterized (simple basis vectors were used). To ensure that each protein concentration model fully encompassed the peak for each component but was not larger than necessary, we performed several trial refinements with REGALS while varying the region of support and inspecting a plot of residual *χ*^2^ vs. frame number (not shown). The model parameters are summarized in Supplementary Table S1. Finally, REGALS was run for 50 iterations, at which point it was well-converged. The overall reduced *χ*^2^ was 1.011, suggesting that the refined model accounted for most of the signal.

The results obtained by REGALS are shown in Figure 2d. The concentrations of the background components (B1 and B2) rise in an approximately linear fashion during elution (Figure 2d, left panel, green and red), reflecting the influence of the smoothness regularizer on these components. The corresponding SAXS profiles show that B1 is associated with an increase in scattering at high-*q*, while B2 is primarily a low-*q* feature (Figure 2d, right panel, green vs. red). The protein components (C1 and C2) have compact peaks with positive concentrations and corresponding SAXS profiles that appear well-subtracted (Figure 2d, blue and orange). We previously showed that these protein components were in excellent agreement with models of the monomeric and dimeric forms derived from crystal structures [31].

Although SVD had suggested that two background components were needed to describe the data, we asked whether two components were strictly necessary for deconvolving the protein peaks. To test this, we removed the minor component from the model (B2) and performed the deconvolution using REGALS. As expected, the quality of fit was noticeably worse when the background was modeled with one component compared with two (Figure 3a, bottom vs. top). Interestingly, the fit of the one-background model is worse in the buffer-only region of the data, but it achieves a near-perfect fit (*χ*^2^ ~ 1) in the region where the proteins elute. This observation suggests that the protein components absorbed the background subtraction error. Indeed, a comparison of the extracted SAXS profiles for C1 shows a significant deviation from expected shape in the low-*q* region if only one background is used (Figure 3b). These results indicate that the buffer scattering in AEX-SAXS can be complex and must be modeled well to achieve well-subtracted SAXS profiles. Furthermore, they underscore the importance of collecting the full buffer scattering before and after the peak in AEX-SAXS experiments, as this information is effectively used to extrapolate the complex behavior underneath the elution peaks.

**Figure 3:**
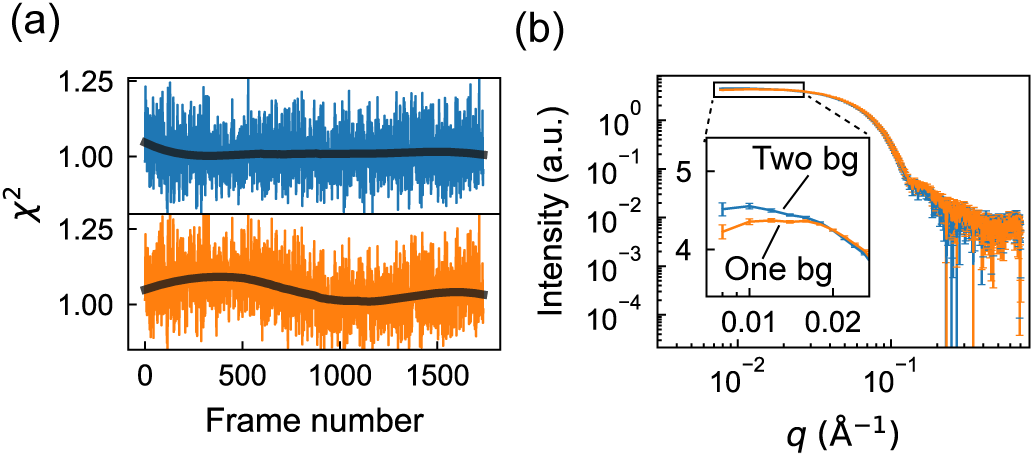
The changing background in AEX-SAXS can be complex. (**a**) Comparison of *χ*^2^ values from the REGALS deconvolution of AEX-SAXS dataset in Figure 2 with two background components (top panel, blue) versus one (bottom panel, orange). The former is relatively uniform around the expected value of 1, whereas the latter shows unevenness throughout the elution (black curves: smoothed *χ*^2^ values shown as trend lines), indicating that dataset is better described with two background components. (**b**) Because the background scattering includes significant low-*q* features, failing to properly account for the changes in the background can lead to artifacts in the extracted protein scattering profiles. Here, the use of only one background component in the analysis leads to a downturn in the low-*q* region of the component 1 scattering, which will lead to underestimation of protein size.

### 3.2 REGALS deconvolution with real-space SAXS restraints

In SAXS datasets from time-resolved or ligand titration experiments, it is common for components to have non-zero concentrations in most or all of the measurements, and a compact support cannot be assumed in the concentration basis, as in the AEX-SAXS example above. To robustly deconvolve such datasets, it is necessary to incorporate additional prior information. Within REGALS, this can be done in two ways: (1) by imposing boundary conditions on the concentration basis vectors, and (2) by limiting the maximum dimension of certain components through real-space parameterization of the SAXS basis vectors.

#### 3.2.1 Equilibrium titrations

As a first test of real-space restraints, we examined a challenging ligand titration dataset of phenylalanine hydroxylase (PheH) [28]. The tetrameric enzyme undergoes a conformational change upon binding its allosterically-activating ligand, L-phenyalanine (L-phe). In SAXS, the signature of this conformational change is an oscillating mid-*q* feature that appears at physiological concentrations of L-phe (Figure 4a, 0-1 mM L-phe). At higher concentrations of ligand, the mid-*q* feature does not change further, but an increase in scattering at low-*q* is observed (Figure 4a, 3-80 mM L-phe), indicating an increase in the average molecular weight. This larger oligomer or aggregate is likely non-physiological, and therefore previous analysis focused on the 0-1 mM concentration range. However, SEC-SAXS experiments at 0 and 1 mM L-phe revealed the presence of a small amount of aggregate, indicating that all of the SAXS curves in the titration were corrupted by aggregation to some extent, inflating estimates of size and molecular weight even at low L-phe concentrations. The presence of a small population of aggregates is extremely common in SAXS, and thus, a direct method to deconvolve it from other components is of particular interest.

**Figure 4:**
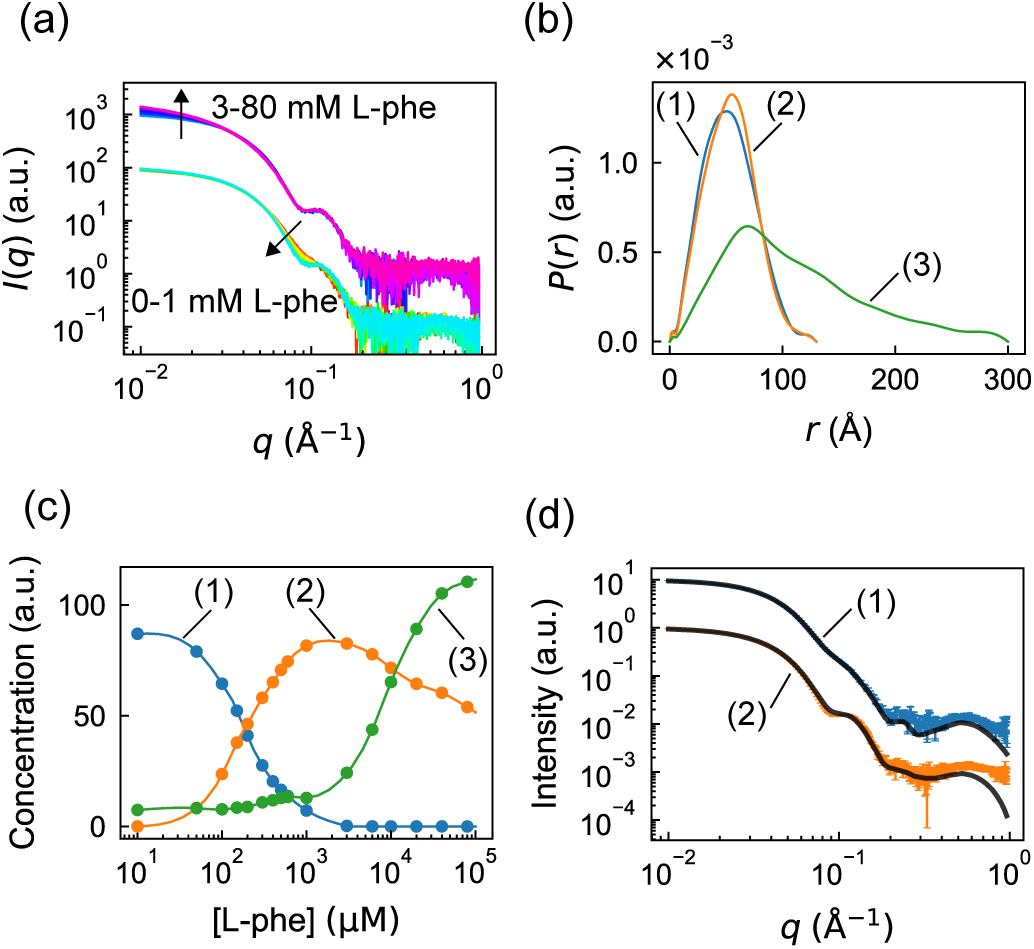
Separation of aggregation from ligand-induced conformational changes in a titration dataset with real space regularization in REGALS. (**a**) Scattering profiles from a previously reported phenylalanine (L-phe) titration experiment on PheH [28]. Up to 1 mM L-phe (red to cyan), the change in scattering mainly occurs at mid-*q*, corresponding to internal conformational changes. At [L-phe] greater than 1 mM (blue to magenta), an increase at low-*q* can be observed, indicative of aggregation. The two sets of profiles are offset for clarity. (**b**) Regularized *P*(*r*) functions from REGALS deconvolution. Different cutoffs for *P*(*r*) functions differentiate aggregation (green) from normal conformations (blue and orange). (**c**) Concentration profiles from REGALS deconvolution are consistent with observation from panel **a**, with conformational switching occurring below 1 mM L-phe and aggregation gradually becoming dominant above 1 mM L-phe. (**d**) Extracted profiles for components 1 and 2 agree with the scattering profiles of inactive and activated PheH, respectively. Here, the black curves are the *P*(*r*) regularized scattering profiles from SEC-SAXS [28].

To deconvolve the PheH titration dataset, we constructed a REGALS model with three components: resting tetramer, activated tetramer, and aggregate. The SAXS profiles for each component were modeled using the real-space parameterization (*P*(*r*)). The resting and activated tetramers were estimated to have a maximum dimension of 130Å based on previous studies [28], and the aggregate was assigned a maximum dimension of 300 Å, the largest dimension that could be measured based on the Shannon limit for this dataset (*d*_max_ < π/*q*_min_). Boundary conditions of *P*(*r*) = 0 were imposed at both *r* = 0 and r = *d*_max_. We also imposed prior information on the concentration basis vectors using a smooth parameterization. According to the equilibrium model for this system [28], the concentration of activated tetramer is negligible at 0 mM L-phe, so a zero boundary condition was imposed. For the resting tetramer, we limited the range of the basis vector to 0-3 mM and imposed a zero boundary condition at 3 mM based on the observation that the mid-*q* feature saturates above this concentration. No limits or boundary conditions were imposed on the aggregate concentration. The independent variable x was calculated as the logarithm of [L-phe], reflecting the higher density of samples at low [L-phe] and the standard practice of visualizing titration data on a logarithmic scale. Regularization was used to enforce smoothness of the concentration profiles and the *P*(*r*) functions. The model parameters are summarized in Supplementary Table S2. The basis vectors were optimized using the REGALS algorithm, which converged after 50 iterations with an overall reduced *χ*^2^ of 1.41.

One advantage of using the real-space parameterization is that *P*(*r*) functions are obtained directly from the deconvolution and provide immediate insight into particle shape. For the resting and activated PheH tetramers, we find that the *P*(*r*) functions decay to zero smoothly at *d*_max_ (Figure 4b, components 1 and 2), as expected for compact particles. The peak in *P*(*r*) shifts toward larger dimensions in the activated tetramer, indicating that it has a less compact conformation. The *P*(*r*) for the aggregated species decays toward zero at *d*_max_ in an approximately linear fashion, which is characteristic of elongated or rod-like shapes (Figure 4b, components 3).

In ligand titration datasets like this one, the concentrations of different components are often of great interest, since they give insight into the equilibrium behavior of the system, including cooperativity and binding affinities. The REGALS deconvolution of the PheH titration produced concentration profiles that appear physically reasonable (Figure 4c). To verify that smoothness regularization had not overly biased the result, we also extracted concentration estimates at each point without regularization (Equation 27) and found that they agree with the regularized curve (Figure 4c, circles vs. continuous curves). We find that the aggregate is present under all conditions, staying at a low level between 0 and 1 mM L-phe, before rising sharply at high concentrations, in agreement with prior SEC-SAXS experiments [28]. The resting tetramer converts into the activated tetramer in a manner characteristic of cooperative two-state transition, as shown previously [28].

Further analysis of this equilibrium is beyond the scope of this study. However, we note that the arbitrary concentration scale of Figure 4c can be transformed readily into the fraction of resting and activated species, which can be fit using an equilibrium model. The REGALS results are normalized by the area under *P*(*r*) (equal to *I*(*q* → 0)), and this quantity is expected to be the same for components with the same molecular weight. Thus, in this case the tetramer concentrations in Figure 4c differ from the true concentrations (e.g. in mg/mL) by the same scale factor.

One assumption in the REGALS model was that the aggregate did not change in size or shape as a function of [L-phe], which may not be the case, particularly since its shape appears to be rod-like and therefore its growth might be non-terminating. To check whether this assumption was supported by the data, we examined the reduced *χ*^2^ of the model at each [L-phe] concentration. Interestingly, *χ*^2^ at the highest [L-phe] concentration is 3.4, which is significantly larger than at other concentrations (1.3 on average). Thus, it seems likely that the scattering profile of the aggregate does change, at least at very high L-phe concentrations. If this is the case and the aggregate was improperly modeled by REGALS, another technique such as SEC-SAXS might be necessary to obtain reliable scattering curves for the tetramers. Nonetheless, the tetramer SAXS curves extracted from the REGALS deconvolution are in excellent agreement with those obtained by SEC-SAXS (Figure 4d), suggesting that inaccuracy of the aggregation model had a minimal effect on deconvolution.

#### 3.2.2 Time-resolved SAXS

Based on the successful application of real-space REGALS to the challenging PheH titration dataset, we asked whether similar models might be applied to time-resolved SAXS. Time-resolved experiments can be performed with two different techniques: mixing and pump-probe. In mixing experiments, a rapid change in solution conditions (such as by rapid dilution, addition of denaturant, allosteric ligand, or reactant) is followed by SAXS measurements after some time has elapsed. This technique is well-suited to irreversible reactions or those that cannot be initiated except by mixing. In contrast, pump-probe experiments are usually initiated by a laser pulse, and followed after a time delay by the X-ray measurement. Compared with mixing, pump-probe measurements can access very short time scales if fast lasers and pulsed X-ray sources are used. Pump-probe datasets are also special in that very small changes can be measured by examining difference profiles (laser on minus laser off), which removes systematic error. Given these differences, we chose to evaluate REGALS with both mixing and pump-probe datasets, as described below.

First, we chose to analyze a stopped-flow mixing dataset from the soluble nucleotide binding domains (NBDs) of the membrane transport protein MsbA, which was recently published and deposited in a public database [39]. In the experiment, a solution with nucleotide-free NBD monomers was rapidly mixed with ATP, resulting in ligand binding and dimerization, followed by ATP hydrolysis and dissociation back to the monomeric state. This transient increase in average size can be observed in a Kratky plot (*q*^2^*I*(*q*) vs. *q*), where the main peak shifts to the left (lower *q*) and then to the right (Figure 5a). In the original publication, the relative concentrations of NBD monomer and dimer at each time point were fit using calculated scattering profiles from known crystal structures. However, in time-resolved experiments generally, it is often the case that atomically detailed structures are not available, either because they have not been characterized at high resolution or because they are dynamic. Therefore, we asked whether REGALS could deconvolve the MsbA dataset using only general properties of the molecules.

**Figure 5:**
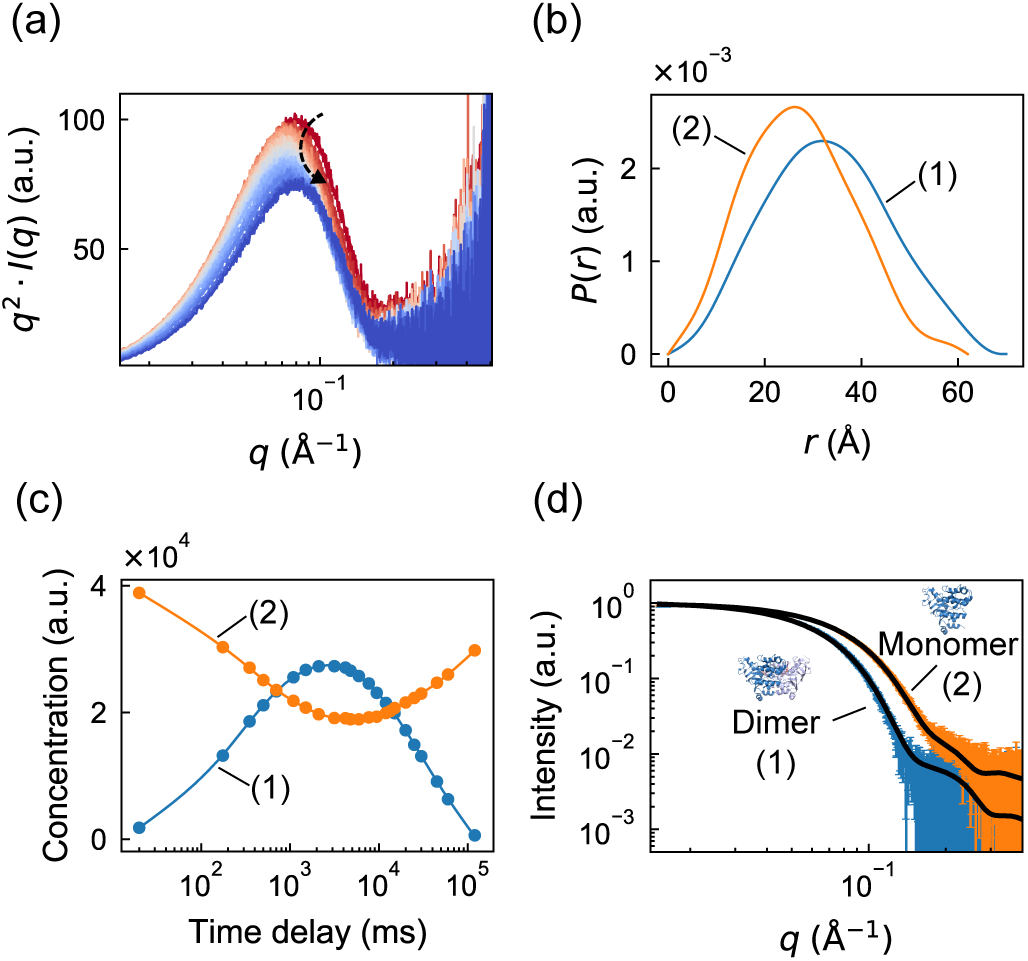
Model-free deconvolution of a time-resolved mixing dataset in REGALS. (**a**) Scattering profiles from a previously reported time-resolved mixing experiment with MsbA NBD and ATP [39] shown as Kratky plots (red to blue). The peak position shifts to *q* ~ 0.07Å^−1^ before returning to *q* ~ 0.08Å^−1^ (denoted by curved arrow), indicating a transient increase in size. (**b**) Regularized *P*(*r*) functions of dimer (blue) and monomer (orange) components have well-defined shapes with *d*_max_ estimates based on the crystal structure of full-length dimeric MsbA (PDB: 3B60). (**c**) Concentration profiles from REGALS deconvolution show transient formation of the NBD dimer. (**d**) The extracted scattering profiles of components 1 and 2 agree with predictions using the NBD dimer and monomer from the full-length crystal structure (black curves: CRYSOL fits).

The REGALS model consisted of two components representing NBD monomer and dimer. The parameterization was similar to the PheH titration example above: the smooth parameterization was used for concentrations, and real-space for SAXS profiles. Based on the full-length structure of dimeric MsbA (PDB ID: 3b60), we estimated the *d*_max_ of the dimeric and monomeric forms of the NBD portions to be 70Å and 62 Å respectively. Reflecting the prior observation that NBDs relax to a fully dimeric state after ATP hydrolysis, we applied a zero boundary condition to the dimer concentration at the final time point (approximately 2 minutes after mixing). The model parameters are summarized in Supplementary Table S3. REGALS was run for 100 iterations, resulting in an overall reduced *χ*^2^ of 0.335. The fact that *χ*^2^ < 1 here suggests that the reported experimental errors were overestimated, so *χ*^2^ is not a reliable statistic for quality of fit. However, the quality of fit was confirmed by examining the residual (not shown).

The deconvolved concentration profiles show a rise and fall of the dimer component, with a concomitant dip in the monomer, which resembles the profiles obtained by fitting scattering from crystal structure models in the original publication [39]. The *P*(*r*) functions are also physically reasonable, with single peaks that decay smoothly to zero as r approaches the maximum dimension. Using the REGALS deconvolution, we extracted the SAXS profiles (Equation 27 in Methods) for the monomer and dimer and compared them with models derived from crystallography (Figure 5d). The excellent agreement suggests that the atomistic models accurately reflect the structures of the NBDs in solution. Although the analysis presented here used estimates for the maximum dimension based on a crystal structure, no assumptions were made about the shape of the individual components.

For a pump-probe dataset, we chose a temperature-jump SAXS/WAXS experiment which was performed on the protein CypA [40]. These experiments involved rapidly heating the sample by approximately 10 °C with an infrared laser pulse of several nanoseconds duration, followed by a synchrotron X-ray pulse of approximately 500 ns duration after a delay of 562 ns to 1 ms. Following the methods in the original publication [40], difference profiles were constructed (laser on minus laser off) for both the protein and buffer blanks, and these were scaled together in the WAXS regime and subtracted. The remaining signal, attributed to the effect of the rapid temperature change, is most significant in the SAXS regime (Figure 6a), and it evolves non-trivially as a function of the time delay (Figure 6a, inset). Note that the difference profiles are negative, and this is thought to result from the differential thermal expansion coefficients of protein and water, which would reduce scattering contrast at high temperature [40].

**Figure 6:**
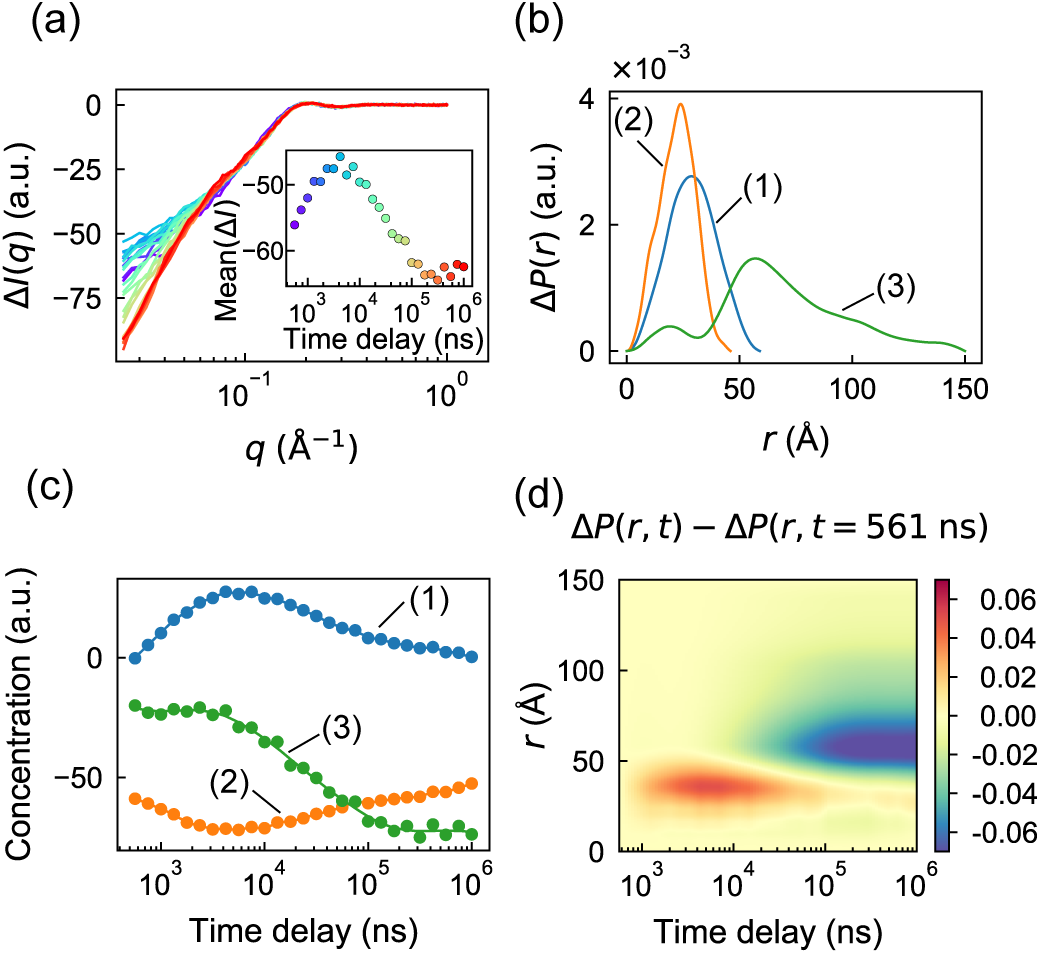
Separation of changes at different length scales in a time-resolved T-jump dataset in REGALS. (**a**) Difference scattering from a previously reported time-resolved T-jump experiment on CypA [40] as a function of time delay (violet to red). Inset: the mean intensity Δ*I* over *q* = 0.03 − 0.05A increases before decreasing. (**b**) Regularized Δ*P*(*r*) functions from REGALS deconvolution. Three cutoffs were chosen to separate change at different length scales: the equilibrium CypA structure (49 Å, orange), thermally-excited intermediate (59 Å, blue), and large length scale change (150 Å, green). (**c**) Concentration profiles from REGALS deconvolution show different kinetics for conformational changes at different length scales. (**d**) Reconstructed Δ*P*(*r, t*) from deconvolved components display two distinct processes occurring at small and large length scales.

Previously, the biphasic appearance of the mean intensity (Figure 6a, inset) was interpreted as a fast transition to excited states of the molecule followed by a slow relaxation toward equilibrium [40]. Although SVD analysis revealed 3 significant components, no kinetic or structural interpretation of the basis vectors was reported. We asked whether a real-space REGALS deconvolution might offer additional insight. Based on the SVD result, we chose to model three components (C1, C2, and C3). For all three, a smooth parameterization was used for the concentration basis, and real-space for the SAXS profile basis. The first component (C1) was assigned to represent the transient process following T-jump, with a concentration of zero at both end points. No constraints were applied to the concentrations of the other two components. In real space, C2 was assigned a maximum dimension of 46Å estimated from a crystal structure of CypA (PDB ID: 3k0n). Lacking further information with which to restrain the model, the maximum dimensions for C1 and C3 were adjusted by trial and error based on quality of fit and subjective appearance of the *P*(*r*) functions. The final model parameters are summarized in Supplementary Table S4. Note that since difference intensities are fit, this parameterization represents the difference *P*(*r*) function, Δ*P* = *P*_on_ − *P*_off_, and *d*_max_ represents the maximum dimension over which changes to *P*(*r*) occur after heating.

The REGALS algorithm was run for 400 iterations, converging to an overall reduced *χ*^2^ of 1.667. Although the difference intensities are negative (Figure 6a), the deconvolved Δ*P*(*r*) functions are all positive (Figure 6b) because the REGALS algorithm normalizes SAXS basis functions by the integral of *P*(*r*). Consequently, some of the concentrations are negative (Figure 6c). Negative concentrations (or SAXS curves) are a necessary feature when analyzing difference intensities, and they can be a challenge to conceptualize. However, two immediate observations can be made. First, the concentration of C3 is approximately constant for the first ~ 4 *μ*s after T-jump, and the change on those timescales is captured by C1 and C2. According to the Δ*P*(*r*) functions for C1 and C2, we conclude that the fast processes occur on length scales up to 60Å, somewhat larger than the size of the CypA monomer. Changes on longer timescales additionally involve C3, which has a much longer range of 150Å and likely involve interparticle interactions because the experiments were done at a relatively high protein concentration of 50 mg/mL.

To gain a more intuitive picture of the changes following T-jump, we used the regularized basis functions to reconstruct the time evolution of Δ*P*(*r*). This removes, to some extent, the influence of choices made during the REGALS parameterization, and resolves the sign ambiguity. Since the signal is dominated by the contrast decrease (not shown), we subtracted the first time point to obtain ΔΔ*P*(*r, t*) ≡ Δ*P*(*r, t*) − Δ*P*(*r, t* = 561ns), which tracks the change in signal after T-jump (Figure 6d). This reconstructed signal reveals a clear positive feature with a peak at *r* ~ 35Å that appears at fast time scales, followed by a negative feature with a peak at *r* ~ 60Å on slower time scales. The physical explanation is not entirely clear, but one hypothesis might be transient partial unfolding followed by an increase in inter-particle repulsion (or a decrease in attraction). As experiments which rely on difference intensities are often performed at high protein concentrations, further investigations of inter-particle interactions are of great interest.

## 4 Conclusions

In this work, we introduced REGALS as a robust, generally applicable technique to deconvolve challenging SAXS datasets from evolving mixtures. The strategy implemented in REGALS has several key advantages. Most notably, prior knowledge is taken into account without having to impose a physicochemical “hard” model or known scattering curves. Having flexible restraints is important in cases where such models are not available, or when SAXS is to be used for cross-validation. Second, the method is readily adapted to a range of experiments. As we demonstrated, AEX-SAXS, ligand titrations, time-resolved mixing, and time-resolved pump-probe datasets can all be analyzed successfully by REGALS. Finally, REGALS is not a black box; the model assumptions are physically motivated, easily explained, and completely specified by the user. Because deconvolution can be ambiguous and strongly influenced by model assumptions, this transparency is essential when communicating scientific results.

The flexibility of the REGALS method is reflected in our software implementation (see Methods). The model is specified using object-oriented code, which facilitates mixing and matching parameterizations to suit the experiment. In order to provide feedback to the user and support customization, the code is run using a live notebook that performs data import, model definition, optimization, and visualization. Example notebooks are provided for each of the datasets described here. Since SAXS is a rapidly-developing technique, we designed REGALS with future changes in mind. Its hierarchical object structure allows for new linear parameterizations and quadratic regularizers to be added with minimal changes to the existing code. Finally, to facilitate future development and adoption by the community, we provided two functionally equivalent implementations of REGALS in MATLAB and python. The code is version-controlled, open source, and free to use.

Future work will focus on augmenting the REGALS toolkit to further expand the range of applications. Here, we found that two simple restraints, smoothness and compact support, proved powerful for expressing prior knowledge. However, many other types of restraint are possible within the REGALS framework. Examples of particular interest to SAXS include sparseness and non-negativity [22], hard restraints on certain components with known scattering curves, and fixed non-zero or derivative boundary conditions. In addition, REGALS could be applied to datasets with more than one independent variable using methods from MCR of multi-way data [16]. For example, the CypA time series analyzed here was one among several conducted at different initial temperatures [40], and thus, the entire dataset might be analyzed using a multiway REGALS decomposition. Furthermore, the assumption of dilute solution can also be relaxed by adding extra components to represent terms in the Taylor expansion of the structure factor [25], which may be of particular interest for time-resolved experiments that require high protein concentrations. Finally, certain parameter choices in REGALS may be automated by leveraging the Bayesian interpretation of regularized linear regression [41, 42], much as the regularization parameter and *d*_max_ are determined automatically in Bayesian IFT [36]. We anticipate that the REGALS method described here, and future developments, will be a valuable addition to the SAXS data analysis toolset and enable new applications.

## 5 Methods

### 5.1 Computational details

#### 5.1.1 Least-squares optimization in each REGALS iteration

The two regularized linear least squares problems within each REGALS iteration (Equations 17 and 18) are solved using the method of normal equations. For the SAXS profile basis (Equation 17), the best-fit parameters satisfy *K* sets of linear equations (with *k* = 1,2,…, *K*):

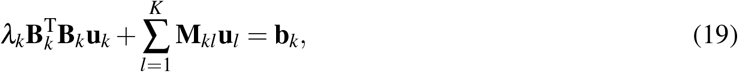

where

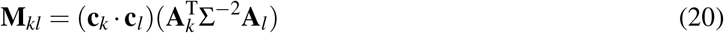

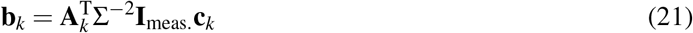

Note that these equations can be combined and written in the form (**M** + **H**)**u** = **b**, making them straightforward to solve using standard numerical methods. Similarly, the parameters for the concentration basis (Equation 18) are found by solving the *K* sets of linear equations:

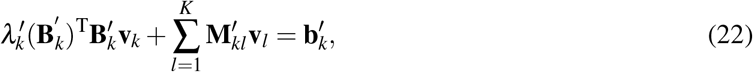

where

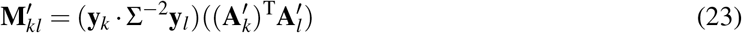

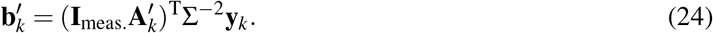

#### 5.1.2 Extracting scattering curves and error estimates

After fitting a dataset with a REGALS model for **Y** and **C**, the results are typically smooth versions of the concentrations and SAXS profiles. However, for further analysis (such as fitting atomistic models to the SAXS data), it is desirable to extract curves resembling experimental data with properly estimated errors. Previously, we applied a projection algorithm which uses the pseudoinverse of the concentration matrix to generate SAXS profiles and associated error bars [28]. For the datasets examined here, we found that the pseudoinverse method amplifies noise in certain cases. Therefore, we developed an alternate method which makes use of the regularized basis vectors to overcome this issue. In order to extract a particular component, a residual data matrix is reconstructed by subtracting the model with component *k* excluded:

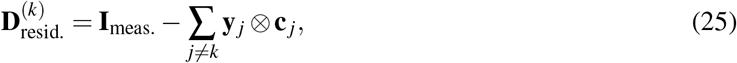

The unregularized basis functions **y** and **c** are extracted by minimizing

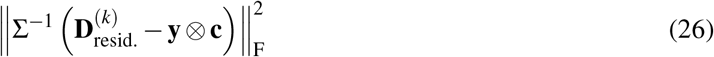

with either scattering profile or concentration held fixed. The solutions can be written as weighted averages of the residual data matrix, as follows:

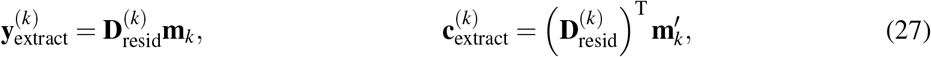

with coefficients

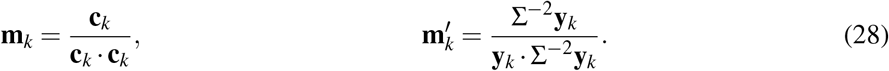

The uncertainties are estimated by standard propagation of experimental errors:

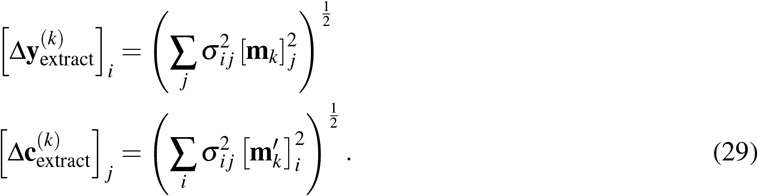

#### 5.1.3 Regularization parameter estimation

The regularization parameters *λ_k_* and 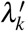 reflect prior information about the smoothness of the parameters. They are not known in advance, so initial values must be chosen by the user and further adjusted if REGALS fails to converge. However, the regularization parameter is not an intuitive quantity, and it depends in a complicated fashion on the noise level in the data and the particular regularizer chosen. To assist the user in selecting initial values, we provide the option of specifying a more intuitive parameter: the “number of good parameters,” or *n_k_*. This parameter comes from the Bayesian interpretation of regularized linear regression [41, 42], and it estimates how many parameters are effectively determined by the data (as opposed to the regularizer). The number of good parameters determined for **u**_*k*_ (SAXS basis) is as follows:

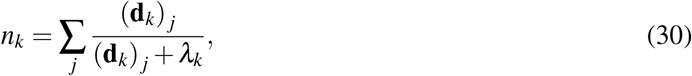

where **d**_*k*_ is the vector of generalized eigenvalues of the matrices **M**_*kk*_ (which depends on |**c**_*k*_|, see Equation 20) and 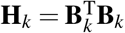. To determine *λ_k_* given *n_k_*, Equation 30 is solved numerically using the initial guess for **c**_*k*_. Strictly speaking, *n_k_* should be determined using the final value of **c**_*k*_ (after REGALS has converged), however we have found that initial estimates of *n_k_* are usually close to the final values. Similarly, regularization parameters for the concentration basis are found by solving Equation 30 where **d**_*k*_ are the generalized eigenvalues of 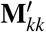 (Equation 23) and 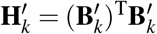.

#### 5.1.4 Software implementation

The REGALS method was developed in MATLAB and subsequently translated into python. The two implementations have similar organization and produce equivalent results. Both versions are available for the convenience of future users and developers.

The code is organized using a hierarchy of classes to facilitate mixing-and-matching basis vector types (Figure 1b). At the lowest level are Concentration and Profile classes for each type (simple, smooth, and real-space), which share a common interface and are responsible for calculating the **A**_*k*_ and **B**_*k*_ matrices (Equations 13 and 16) given parameters such as boundary conditions, number of samples and extent. At an intermediate level is the Component class, which represents a single component in the mixture and contains one Concentration object and one Profile object. At the top level is the Mixture class, which contains an array of Component objects as well as the parameter vectors and regularization parameters. Methods are included to compute the terms appearing in the normal equations (Equations 19 and 22), estimate regularization parameters (inversion of Equation 30), and extract basis vectors (Equations 27 and 29). Finally, the REGALS class implements alternating least-squares, and it includes a high-level method (REGALS.run) that controls flow through the algorithm with user-specified termination conditions.

The process of setting up, running, and analyzing a REGALS calculation is performed by writing scripts to interact with the objects. We have included example scripts in the form of live notebooks (Jupyter notebooks in Python) for each of the application examples presented here. Source code, documentation, and examples are available at https://github.com/ando-lab/regals. The release associated with this publication has been tagged as v1.0.

### 5.2 Example data

#### 5.2.1 AEX-SAXS of BsRNR large subunit

The collection and preprocessing of AEX-SAXS from the large subunit of B. subtilis ribonucleotide reductase (BsRNR) was described in the original publication [31]. Briefly, the as-isolated protein was eluted from a MonoQ column using a linear gradient of 100 to 500 mM NaCl directly into a SAXS flow cell. Scattering images were recorded continously during elution (q-range of 0.008 to 0.700 Å^−1^). After integration, each profile was normalized by the transmitted beam intensity, and buffer-only curves collected before the start of the gradient were averaged and subtracted from the remaining curves. A set of 1737 frames was retained for further analysis, beginning just after the start of the linear gradient and ending before the gradient completed, when the NaCl concentration had reached approximately 400 mM. These preprocessed data are available in NrdE_mix_AEX.mat (a MATLAB-formatted hdf5 file).

#### 5.2.2 Equilibrium titration of PheH

A SAXS titration of phenylalanine hydroxylase (PheH) with phenylalanine (L-phe) was performed previously [28]. In the original publication, SAXS curves from PheH at 25 μM (monomer concentration) were processed to produce 16 background-subtracted scattering curves, each with a different amount of L-phe (0, 0.05, 0.1, 0.15, 0.2, 0.3, 0.4, 0.5, 0.6, 1, 3, 6, 10, 20, 40, and 80 mM). The same amount of L-phe was present in the buffer-only samples used for subtraction. The q-range was 0.01 to 0.96 Å^−1^, and the scattering was normalized by the transmitted beam intensity. These preprocessed data are available in PheH_titration.mat (a MATLAB-formatted hdf5 file).

#### 5.2.3 Time-resolved mixing of MsbA NBD with ATP

As a first example of time-resolved SAXS data, we chose a recently published stopped-flow mixing dataset [39]. In the experiment, a soluble nucleotide binding domain (NBD) construct (residues 330-581 of the ATP-binding cassette transporter MsbA) was mixed with Mg^2+^ - ATP in a 1:1 (v/v) ratio (final concentrations 500 *μ*M NBD and 450 *μ*M ATP). One X-ray exposure of 35 ms was acquired per shot after a variable time delay of 20 ms to 120 s.

The time-resolved MsbA NBD dataset consisting of 23 buffer-subtracted scattering curves (0.01 < q < 0.5Å^−1^) was downloaded from a public database (https://www.sasbdb.org/data/SASDGV5/), minimally reformatted, and saved as MsbA_time_resolved.mat (a MATLAB-formatted hdf5 file). Minor pre-processing was performed before running REGALS. Upon inspection, we noted a strong negative-going feature at low-*q* suggesting a background subtraction error. We therefore truncated the low-*q* at 0.015Å^−1^. In addition, we found that the average intensity displayed a slight random jitter shot-to-shot. We corrected for this by applying a scale factor to each curve, which was found by fitting a 5th-order polynomial to the mean intensity vs. log_10_(time). The resulting scale factors were close to 1 (standard deviation of 0.013).

#### 5.2.4 Time-resolved temperature-jump of CypA

As an example of a pump-probe time-resolved experiment, we chose recently reported T-jump SAXS/WAXS data collected on the cis-proline isomerase CypA [40]. Here we analyze one particular set of experiments corresponding to the wild type CypA protein and buffer blanks following T-jump to 29.9 ± 0.1 °C. After downloading the raw T-jump data ([43]), we repeated the published data reduction protocol [40] using a custom MATLAB script (available upon request). Briefly, difference scattering curves (Δ*I* = *I*_on_ − *I*_off_) were calculated for both the protein and buffer blanks, and a series of scaling operations was performed to correct for shot-to-shot variations, most crucially the scaling of buffer difference profiles in order to minimize Δ*I*_protein_ − Δ*I*_buffer_ in the WAXS regime, where solvent scattering predominates. This produced a set of 28 difference profiles: 27 logarithmically distributed time points after T-jump and one control where the laser was off prior to X-ray exposure. The control profile was close to zero, indicating that the data processing had not introduced errors, and the remaining profiles resembled those reported in the original publication. After initial data reduction, the WAXS data were discarded and the SAXS portion of the curves (0.025 ≤ *q* < 1Å^−1^) was saved as CypA_Tjump.mat (a MATLAB-formatted hdf5 file).

## Supporting information

Supplementary Tables S1-S4

## Acknowledgments

We thank M.B. Watkins and W.C. Thomas for critical reading of the paper. The authors declare no conflicts of interest.

