## Supporting Information for "REGALS: a general method to deconvolve X-ray scattering data from evolving mixtures"

Meisburger *et al.*

### Contents

**Supplementary Tables** **2**

### Supplementary Tables

| | **B1** (salt gradient) | **B2** (salt gradient) | **C1** (dimer) | **C2** (monomer) |
|---|---|---|---|---|
| **Concentration basis** | | | | |
| Model | *smooth* | *smooth* | *smooth* | *smooth* |
| Range ($x_{\min}$–$x_{\max}$)[a] | 1–1737 | 201–1737 | 730–1270 | 1150–1600 |
| Control points | 50 | 50 | 50 | 50 |
| Zero boundary conditions | none | $x_{\min}$ | $x_{\min}, x_{\max}$ | $x_{\min}, x_{\max}$ |
| Regularization parameter ($\lambda'$)[b] | $7.8 \times 10^8$ | $4.5 \times 10^8$ | 0 | 0 |
| **SAXS basis** | | | | |
| Model | *simple* | *simple* | *simple* | *simple* |

[a] $x$ = frame number
[b] Estimated using $n_g = 8$ (see Equation **??**)

**Table S1:** REGALS model for *Bs*RNR AEX-SAXS dataset

| | **C1** (resting tetramer) | **C2** (activated tetramer) | **C3** (unknown oligomer) |
|---|---|---|---|
| **Concentration basis** | | | |
| Model | *smooth* | *smooth* | *smooth* |
| Range ($x_{\min}$–$x_{\max}$)[a] | 1–5 | 1–3.5 | 1–5 |
| Control points | 31 | 21 | 31 |
| Zero boundary conditions | $x_{\min}$ | $x_{\max}$ | none |
| Regularization parameter ($\lambda'$) | 10 | 10 | 10 |
| **SAXS basis** | | | |
| Model | *real-space* | *real-space* | *real-space* |
| $d_{\max}$ (Å) | 130 | 130 | 300 |
| Control points | 101 | 101 | 101 |
| Zero boundary conditions | $r = 0$, $d_{\max}$ | $r = 0$, $d_{\max}$ | $r = 0$, $d_{\max}$ |
| Regularization parameter ($\lambda$) | $1 \times 10^{12}$ | $1 \times 10^{12}$ | $1 \times 10^{12}$ |

[a] $x = \log_{10} c$ for $c > 0$ and $x = 1$ for $c = 0$, where $c$ is the ligand concentration in $\mu$M

**Table S2:** REGALS model for PheH titration dataset

| | C1 (dimer) | C2 (monomer) |
|---|---|---|
| **Concentration basis** | | |
| Model | *smooth* | *smooth* |
| Range ($x_{min}$–$x_{max}$)[a] | 1.3–5.1 | 1.3–5.1 |
| Control points | 31 | 31 |
| Zero boundary conditions | $x_{max}$ | none |
| Regularization parameter ($\lambda'$) | $1 \times 10^{-3}$ | $1 \times 10^{-3}$ |
| **SAXS basis** | | |
| Model | *real-space* | *real-space* |
| $d_{max}$ (Å) | 70 | 62 |
| Control points | 101 | 101 |
| Zero boundary conditions | $r = 0$, $d_{max}$ | $r = 0$, $d_{max}$ |
| Regularization parameter ($\lambda$) | $1 \times 10^{11}$ | $1 \times 10^{11}$ |

[a] $x = \log_{10}(t)$ where $t$ is the delay time in ms

**Table S3:** REGALS model for MsbA NDB time-resolved dataset

| | C1 (transient) | C2 (intraparticle) | C3 (interparticle) |
|---|---|---|---|
| **Concentration basis** | | | |
| Model | *smooth* | *smooth* | *smooth* |
| Range ($x_{min}$–$x_{max}$)[a] | 2.75–6 | 2.75–6 | 2.75–6 |
| Control points | 31 | 31 | 31 |
| Zero boundary conditions | $x_{min}$, $x_{max}$ | none | none |
| Regularization parameter ($\lambda'$) | 10 | 10 | 10 |
| **SAXS basis** | | | |
| Model | *real-space* | *real-space* | *real-space* |
| $d_{max}$ (Å) | 59 | 46 | 150 |
| Control points | 101 | 101 | 101 |
| Zero boundary conditions | $r = 0$, $d_{max}$ | $r = 0$, $d_{max}$ | $r = 0$, $d_{max}$ |
| Regularization parameter ($\lambda$) | $1 \times 10^{11}$ | $1 \times 10^{11}$ | $1 \times 10^{11}$ |

[a] $x = \log_{10}(t)$ where $t$ is the time delay in ns

**Table S4:** REGALS model for CypA T-jump dataset
